# A plausible mechanism for *Drosophila* larva intermittent behavior

**DOI:** 10.1101/2020.09.19.304774

**Authors:** Panagiotis Sakagiannis, Miguel Aguilera, Martin Paul Nawrot

## Abstract

The behavior of many living organisms is not continuous. Rather, activity emerges in bouts that are separated by epochs of rest, a phenomenon known as intermittent behavior. Although intermittency is ubiquitous across phyla, empirical studies are scarce and the underlying neural mechanisms remain unknown. Here we present the first empirical evidence of intermittency during *Drosophila* larva free exploration. We report power-law distributed rest-bout and log-normal distributed activity-bout durations. We show that a stochastic network model can transition between power-law and non-power-law distributed states and we suggest a plausible neural mechanism for the alternating rest and activity in the larva. Finally, we discuss possible implementations in behavioral simulations extending spatial Levy-walk or coupled-oscillator models with temporal intermittency.

---

**T**he search for statistical regularities in animal movement is a predominant focus of motion ecology. Random walks form a broad range of models that assume discrete steps of displacement obeying defined statistical rules and acute reorientations. A Levy walk is a random walk where the displacement lengths and the respective displacement durations are drawn from a heavy-tailed, most often a power-law distribution. When considered in a 2D space reorientation angles are drawn from a uniform distribution. This initial basic Levy walk has been extended to encompass distinct behavioral modes bearing different go/turn parameters, thus termed composite Levy walk. Levy walks have been extensively studied in the context of optimal foraging theory. A Levy walk with a power-law exponent between the limit of ballistic (*α* = 1) and brownian motion (*α* = 3) yields higher search efficiency for foragers with an optimum around *α* = 2 when search targets are patchily or scarcely distributed and detection of a target halts displacement (truncated Levy walk) (1).

Nevertheless, the underlying assumption of non-intermittent flow of movement in Levy walk models complicates the identification of the underlying generative mechanisms as they focus predominantly on reproducing the observed spatial trajectories, neglecting the temporal dynamics of locomotory behavior. Therefore, Bartumeus (2009) stressing the need for a further extension coined the term intermittent random walk, emphasizing the integration of behavioral intermittency in the theoretical study of animal movement (2). Here we aim to contribute to this goal by studying the temporal patterns of intermittency during *Drosophila* larva free exploration in experimental data and in a conceptual model, bearing in mind that power-law like phenomena can arise from a wide range of mechanisms, possibly involving processes of different timescales (1). While our study remains agnostic towards whether foragers really perform Levy walks - a claim still disputed (1) - we suggest that intrinsic motion intermittency should be taken into account and the assumption of no pauses and acute reorientations should be dropped in favor of integrative models encompassing both activity and inactivity.

*Drosophila* larva is a suitable organism for the study of animal exploration patterns and the underlying neural mechanisms. A rich repertoire of available genetic tools allows acute activation, inhibition or even induced death of specific neural components. Crawling in 2D facilitates tracking of unconstrained behavior. Also, fruit flies during this life stage are nearly exclusively concerned with foraging. Therefore a food/odor-deprived environment can be largely considered stimulus-free, devoid of reorientation or pause sensory triggers, while target-detection on contact can be considered certain. Truncated spatial Levy-walk patterns of exploration with exponents ranging from 1.5 to near-optimal 1.96 that hold over at least two orders of magnitude have been previously reported for the *Drosophila* larva. The turning-angle distribution, however, was skewed in favor of small angles and a quasi-uniform distribution was observed only for reorientation events ≥ 50° (3). Moreover, it has been shown that these patterns arise from low-level neural circuitry even in the absence of sensory input or brain-lobe function and have therefore been termed ‘null movement patterns’ (3, 4).

Behavioral intermittency has not been described for the fruitfly larva. Previous empirical studies on adult *Drosophila* intermittent locomotory behavior have concluded that the distribution of durations of rest bouts is power-law while that of activity bouts has been reported to be exponential (5) or power-law (4). Genetic intervention has revealed that dopamine neuron activation affects the activity/rest ratio via modulation of the power-law exponent of the rest bouts, while the distribution of activity bouts remains unaffected. This observation hints towards a neural mechanism that generates the alternating switches between activity and rest where tonic modulatory input from the brain regulates the activity/rest balance according to environmental conditions and possibly homeostatic state.

Here we analyze intermittency in a large experimental dataset and present a conceptual model that generates alternation between rest and activity, capturing empirically observed power-law and non-power-law distributions. We discuss a plausible neural mechanism for the alternation between rest and activity and the regulation of the animal’s activity/rest ratio via modulation of the rest-bout power-law exponent by top-down modulatory input. Our approach seeks to elaborate on the currently prevailing view that these patterns result from intrinsic neural noise (4).

## Materials and Methods

### Experimental dataset

We use a larva-tracking dataset available at the DRYAD repository, previously used for spatial Levy-walk pattern detection (3). The dataset consists of up to one hour long recordings of freely moving larvae tracked as a single point (centroid) in 2D space. We consider three temperature-sensitive shibire^*ts*^ fly mutants allowing for inhibition of mushroom-body (MB247),brain-lobe/SOG (BL) or brain-lobe/SOG/somatosensory (BLsens) neurons and an rpr/hid mutant line inducing temperature-sensitive neuronal death of brain-lobe/SOG/somatosensory (BLsens) neurons. Each mutant expresses a different behavioral phenotype when activated by 32°-33° C temperature. We compare phenotypic behavior to control behavior in non-activated control groups. A reference control group has been formed consisting of all individuals of the four 32°-33° C control groups (Tab. 1).

For the present study recordings longer than 1024 seconds have been selected. Instances where larvae contacted the arena borders were excluded. The raw time series of x,y coordinates have been forward-backward filtered with a first-order butterworth low-pass filter of cutoff frequency 0.1 Hz before computing the velocity. The cutoff frequency was selected as to preserve the plateaus of brief stationary periods while suppressing the signal oscillation due to peristaltic-stride cycles. Velocity values ≥ 2.5 mm/sec have been discarded to account for observed jumps in single-larva trajectories that are probably due to technical issues during tracking. This arbitrary threshold was selected as an upper limit for larvae of length up to 5mm, crawling at a speed of up to 2 strides/sec with a scaled displacement per stride of up to 0.25.

### Bout annotation

In order to designate periods of rest and activity we need to define a suitable threshold *V*_*θ*_ in the velocity distribution as done for the adult fruitfly in (5). We used the density estimation algorithm to locate the first minimum *V*_*θ*_ = 0.085*mm/sec* in the velocity histogram of the reference control group. A rest bout is then defined as a period during which velocity does not exceed *V*_*θ*_. Rest bouts necessarily alternate with periods termed activity bouts. The bout annotation method is exemplified for a single larva track in Fig. 1.

**Fig. 1.**
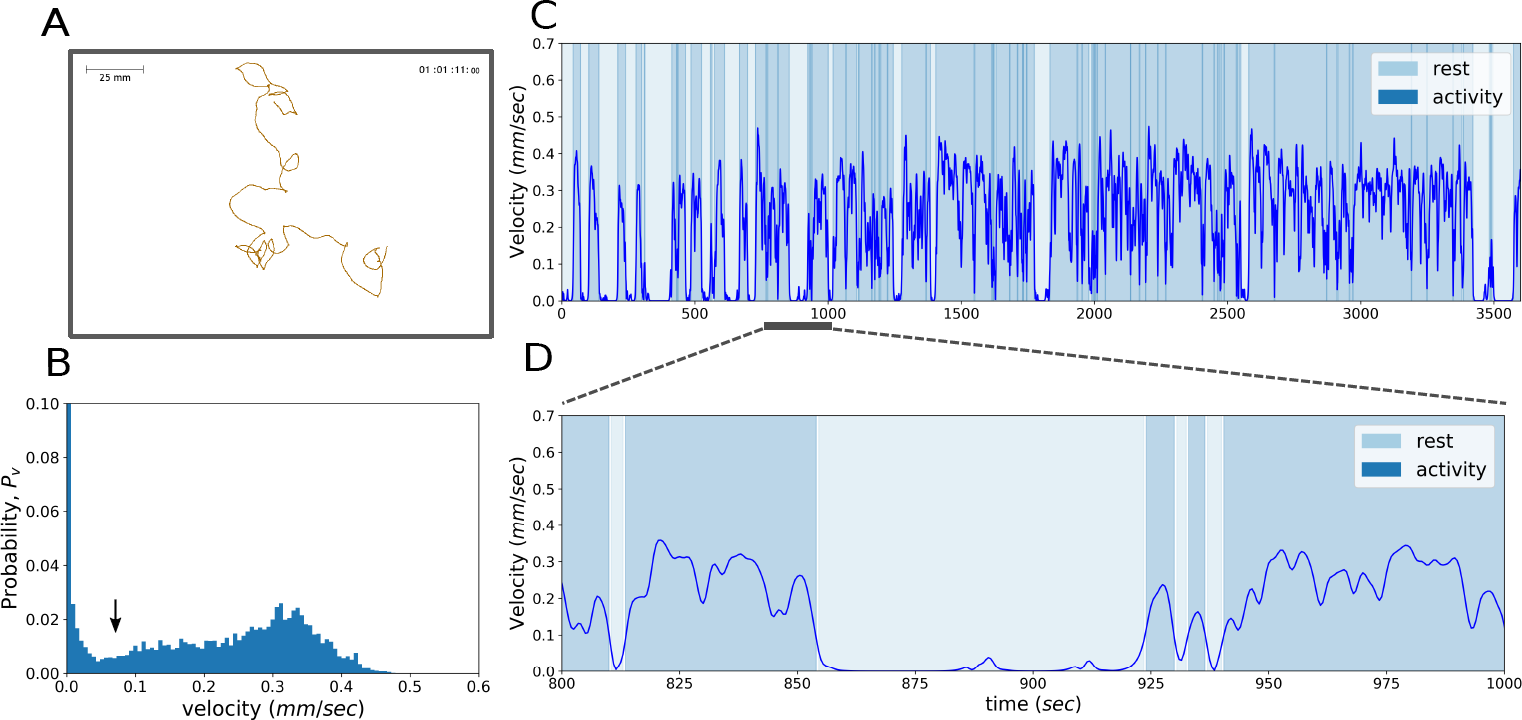
Bout annotation methodology. **A:** Individual larva trajectory. Spatial scale and recording duration are noted. **B:** Velocity distribution for the single larva. The threshold obtained from the reference group, used for rest vs activity bout annotation is denoted by the arrow. **C:** The entire velocity time series of the larva. Rest and activity bouts are indicated by different background colors. **D:** Magnification of the velocity time series.

### Bout distribution

To quantify the duration distribution of the rest and activity bouts we used the maximum likelihood estimation (MLE) method to fit a power-law, an exponential and a log-normal distribution for each group as well as for the reference control group. Given the tracking framerate of 2 Hz and the minimal tracking time of 1024 seconds, we limited our analysis to bouts of duration 2^1^ to 2^10^ seconds. The Kolmogorov-Smirnov distance *D*_*KS*_ for each candidate distribution was then computed over 64 logarithmic bins covering this range. Findings are summarized in Tab. 2 for the rest bouts and in Tab. 3 for the activity bouts.

**Table 1.**
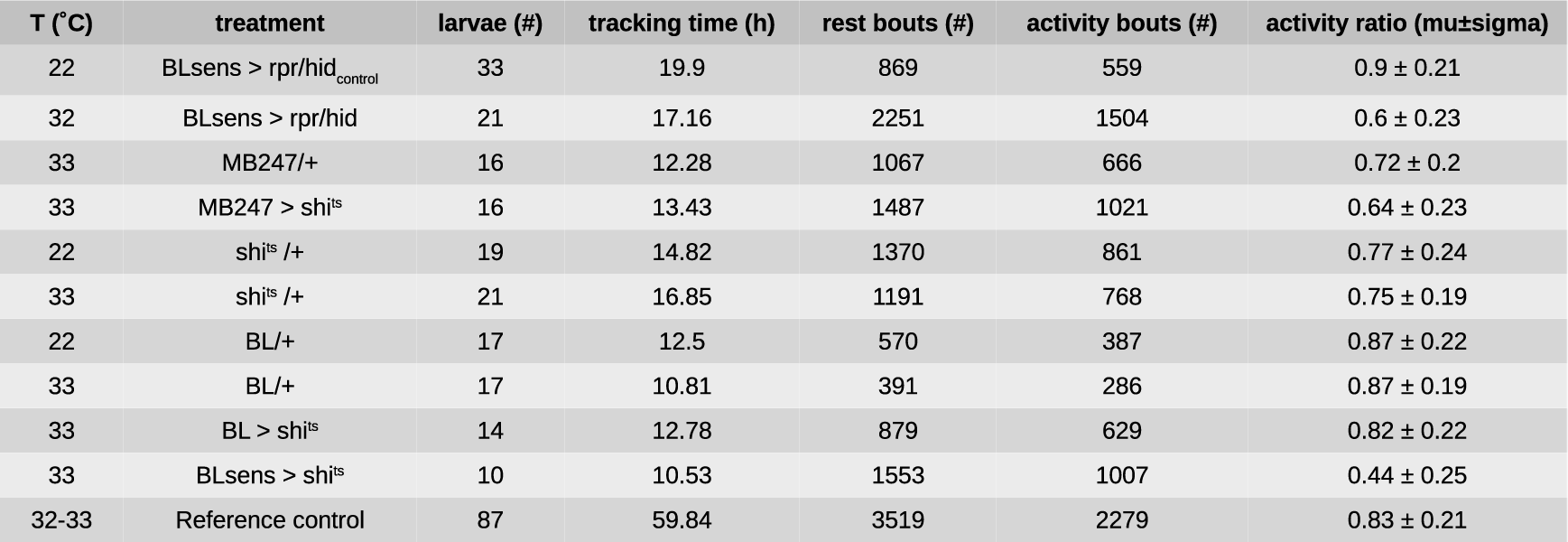
Dataset description and empirical results for rest/activity bout analyses.

**Table 2.** Distribution parameter fits of empirical rest bout duration. The relevant parameters for the best fitting distribution are indicated in bold text.

| T (°C) | treatment | rest bouts |  |  |  |  |  |  |
| --- | --- | --- | --- | --- | --- | --- | --- | --- |
|  |  | powerlaw |  | exponential |  | lognormal |  |  |
|  |  | alpha | KS D | lambda | KS D | mu | sigma | KS D |
| 22 | BLsens > rpr/hid<sub>control</sub> | **1.938** | **0.089** | 0.119 | 0.411 | 1.066 | 1.122 | 0.171 |
| 32 | BLsens > rpr/hid | **1.6** | **0.094** | 0.04 | 0.439 | 1.665 | 1.476 | 0.13 |
| 33 | MB247/+ | **1.58** | **0.06** | 0.042 | 0.438 | 1.724 | 1.55 | 0.133 |
| 33 | MB247 > shi<sup>ts</sup> | 1.41 | 0.167 | 0.029 | 0.246 | **2.438** | **1.574** | **0.073** |
| 22 | shi<sup>ts</sup> /+ | **1.702** | **0.086** | 0.049 | 0.494 | 1.425 | 1.369 | 0.149 |
| 33 | shi<sup>ts</sup> /+ | **1.465** | **0.099** | 0.027 | 0.384 | 2.151 | 1.699 | 0.103 |
| 22 | BL/+ | **1.783** | **0.046** | 0.048 | 0.538 | 1.277 | 1.399 | 0.181 |
| 33 | BL/+ | **1.514** | **0.101** | 0.039 | 0.406 | 1.944 | 1.623 | 0.127 |
| 33 | BL > shi<sup>ts</sup> | 1.666 | 0.109 | 0.111 | 0.246 | **1.502** | **1.189** | **0.103** |
| 33 | BLsens > shi<sup>ts</sup> | 1.483 | 0.125 | 0.033 | 0.387 | **2.072** | **1.554** | **0.105** |
| 32-33 | Reference control | **1.598** | **0.061** | 0.044 | 0.448 | 1.671 | 1.55 | 0.14 |

**Table 3.** Distribution parameter fits of empirical activity bout duration. The relevant parameters for the best fitting distribution are indicated in bold text.

| T (°C) | treatment | activity bouts |  |  |  |  |  |  |
| --- | --- | --- | --- | --- | --- | --- | --- | --- |
|  |  | powerlaw |  | exponential |  | lognormal |  |  |
|  |  | alpha | KS D | lambda | KS D | mu | sigma | KS D |
| 22 | BLsens > rpr/hid<sub>control</sub> | 1.351 | 0.177 | 0.017 | 0.249 | **2.847** | **1.641** | **0.041** |
| 32 | BLsens > rpr/hid | 1.591 | 0.196 | 0.096 | 0.189 | **1.691** | **1.101** | **0.062** |
| 33 | MB247/+ | 1.428 | 0.149 | 0.032 | 0.248 | **2.336** | **1.524** | **0.063** |
| 33 | MB247 > shi<sup>ts</sup> | 1.391 | 0.212 | 0.027 | 0.227 | **2.557** | **1.412** | **0.035** |
| 22 | shi<sup>ts</sup> /+ | 1.371 | 0.175 | 0.025 | 0.212 | **2.699** | **1.528** | **0.066** |
| 33 | shi<sup>ts</sup> /+ | 1.359 | 0.184 | 0.018 | 0.287 | **2.786** | **1.634** | **0.044** |
| 22 | BL/+ | 1.335 | 0.191 | 0.018 | 0.214 | **2.988** | **1.644** | **0.079** |
| 33 | BL/+ | 1.327 | 0.215 | 0.015 | 0.224 | **3.058** | **1.566** | **0.042** |
| 33 | BL > shi<sup>ts</sup> | 1.483 | 0.179 | 0.045 | 0.252 | **2.07** | **1.353** | **0.063** |
| 33 | BLsens > shi<sup>ts</sup> | 1.682 | 0.209 | 0.147 | 0.201 | **1.466** | **1.011** | **0.096** |
| 32-33 | Reference control | 1.366 | 0.173 | 0.019 | 0.26 | **2.73** | **1.617** | **0.046** |

## Results

The results section is organized as follows. Initially we present a simple conceptual two-state model transitioning autonomously between power-law and non-power-law regimes. Next we analyse intermittency during larva free exploration in a freely available dataset (3). Finally we compare mutant and control larva phenotypes in the context of intermittency.

### Network model of binary units reproduces larval statistics of intermittent behavior

Previous work on *Drosophila* adult intermittent behavior reported that rest-bout durations are power-law distributed while activity-bout durations are exponentially distributed (5). Our first contribution is to provide a simple model displaying how this dual regime might emerge. We define a kinetic Ising model with *N* = 1000 binary neurons, with homogeneous all-to-all connectivity (Fig. 2A). Each neuron *i* is a stochastic variable with value *s*_*i*_(*t*) at time *t* that can be either 1 or 0 (active or inactive). We assume that this neuron population inhibits locomotory behavior, so that when Σ_*i*_*s*_*i*_(*t*) > 0 the larva is in the rest phase, and otherwise the larva remains active.

**Fig. 2.**
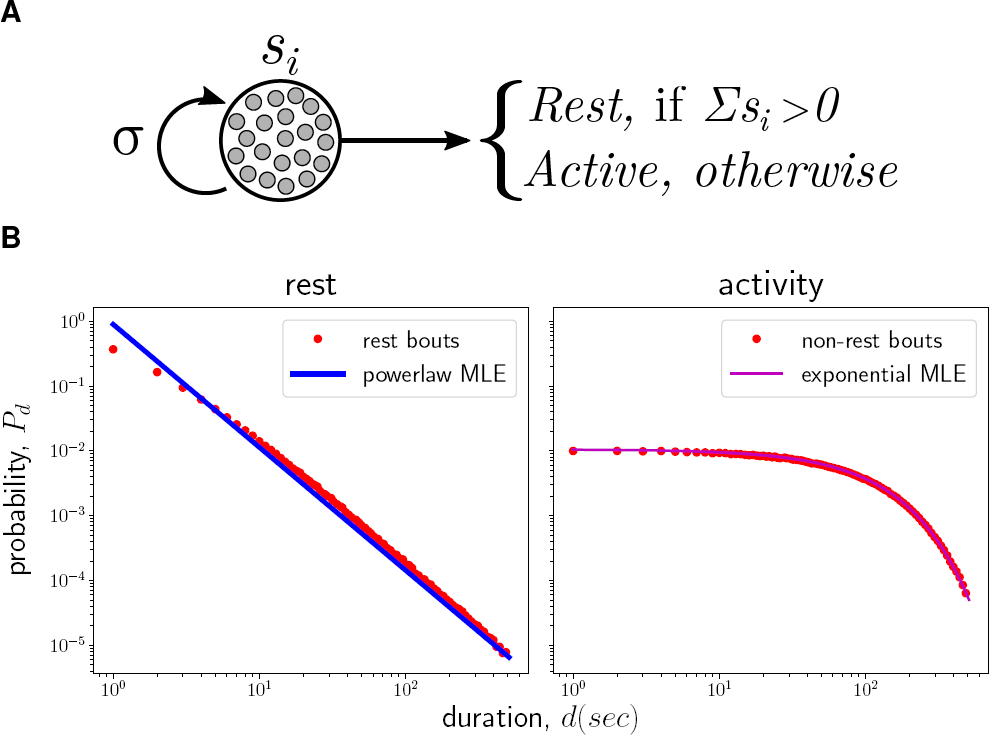
Probability distribution of the duration *d* of rest and activity phases in a branching process model of *σ* = 1, simulated over 10^5^ occurrences of each phase. Duration is measured as the number of updates until a phase is ended. Unit activation *s*_*i*_(*t*) propagates to neighbouring units creating self-limiting avalanches. In the rest phase, when Σ_*i*_*s*_*i*_(*t*) > 0, the system yields a power law distribution with exponent *α ≈* 2. In the activity phase, when Σ_*i*_ *s*_*i*_(*t*) = 0, one unit of the system is activated with probability *µ* = 0.01, yielding an exponential distribution with coefficient *λ* = 0.1.

At time *t* + 1, each neuron’s activation rate is proportional to the sum of activities at time *t*, and will be activated with a linear probability function 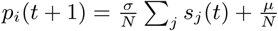. Here, *σ* is the propagation rate, which indicates that when a node is active at time *t*, it propagates its activation at time *t* + 1 on average to *σ* other neurons. When one neuron is activated, this model behaves like a branching process (6), with *σ* as the branching parameter. If *σ* < 1, activity tends to decrease rapidly until all units are inactive while, if *σ* > 1, activity tends to be amplified until saturation. At the critical point, *σ* = 1, activity is propagated in scale-free avalanches, in which duration *d* of an avalanche once initiated follows a powerlaw distribution *P* (*d*) ∼ *d*^*−α*^ (Fig. 2B, left), governed by a critical exponent (*α* = 2 at the *N* → ∞ limit) describing how avalanches at many different scales are generated.

When an avalanche is extinguished, the system returns to quiescence which is only broken by the initiation of a new avalanche. With a residual rate *µ* = 0.01 the system becomes active by firing one unit and initiating a new avalanche. In this case the duration of quiescence bouts (the interval between two consecutive avalanches) follows an exponential distribution (Fig. 2B, right).

This simple conceptual model alternates autonomously between avalanches of power-law distributed durations and quiescence intervals of exponentially distributed durations. This alternation between power-law and non-power-law regimes can serve as a basic qualitative model of the transition between rest and activity bouts in the larva (cf. Discussion).

### Parameterization of larval intermittent behavior

We analyzed intermittent behavior during larval crawling in a stimulus-free environment (cf. Materials and Methods for dataset description). Each individual larva was video-tracked in space (Fig. 1A). From the time series of spatial coordinates we computed the instantaneous velocity and determined a threshold value (Fig. 1B) that separates plateaus of continued activity (activity bouts) from epochs of inactivity (rest bouts, Fig. 1C-D) following the analyses suggested in (5).

We start out with the analysis of experimental control groups that were not subjected to genetic intervention. As a first step we computed the number of occurrences of rest and activity bouts and the activity ratio, which quantifies the accumulated activity time as fraction of the total time (Tab. 1). For the reference control group we obtain an activity ratio of 0.83 albeit with a fairly large variance across individuals.

For the duration distribution of rest bouts we find that it is best approximated by a power-law distribution in all six control groups (Tab. 2) in line with previous results reported for the adult fruitfly (4, 5). The empirical duration distribution of rest-bouts across the reference control group is depicted in Fig. 3 A (red dots). Again, the power law provides the best distribution fit. The exponent *α* of the power law ranges from 1.514 to 1.938 with *α* = 1.598 for the reference control group.

**Fig. 3.**
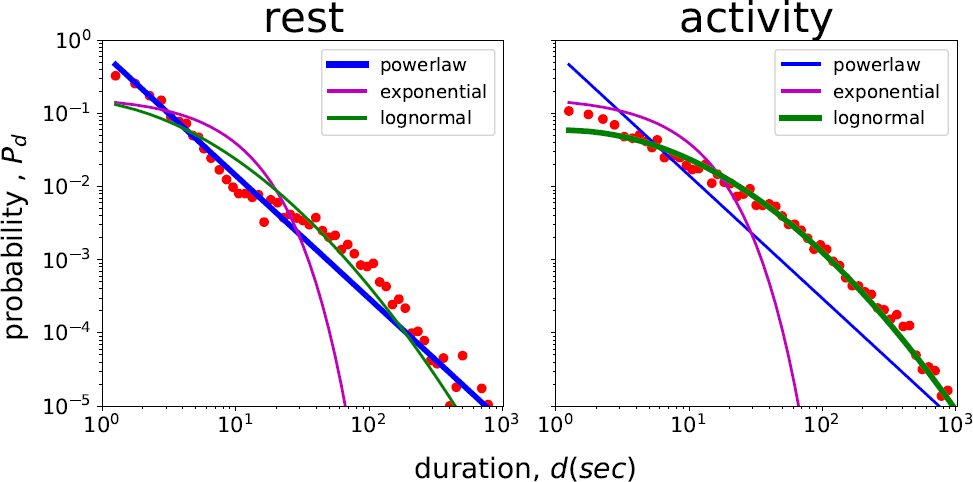
Probability density of rest and activity bout durations for the reference control group. Dots describe the probability density over logarithmic bins. Lines are the best fitting power-law, exponential and log-normal distributions. The thick line denotes the distribution having the minimum Kolmogorov-Smirnov distance *D*_*KS*_ (Tab. 2 - 3).

When analyzing the durations of activity bouts we found that these are best approximated by a log-normal distribution in all groups (Tab. 3). This result is surprising as previous work in the adult suggested the mode of an exponential distribution (5). For the reference control group Fig. 3 B compares the empirical duration distribution of activity bouts with the fits of the three distribution functions.

### Modification of rest and activity bout durations in mutant flies

Behavioral phenotypes in genetic mutants can help identify brain neuropiles in the nervous system of *Drosophila* larva that are involved in the generation of intermittent behavior, or that have an effect on its modulation. To this end we analyzed 4 experimental groups where genetic intervention was controlled by temperature either via the temperature-sensitive shibire protocol or via temperature-induced neuronal death (rpr/hid genotype). Each group is compared to a non-activated control group as shown in Fig. 4 and described in Tab. 1.

**Fig. 4.**
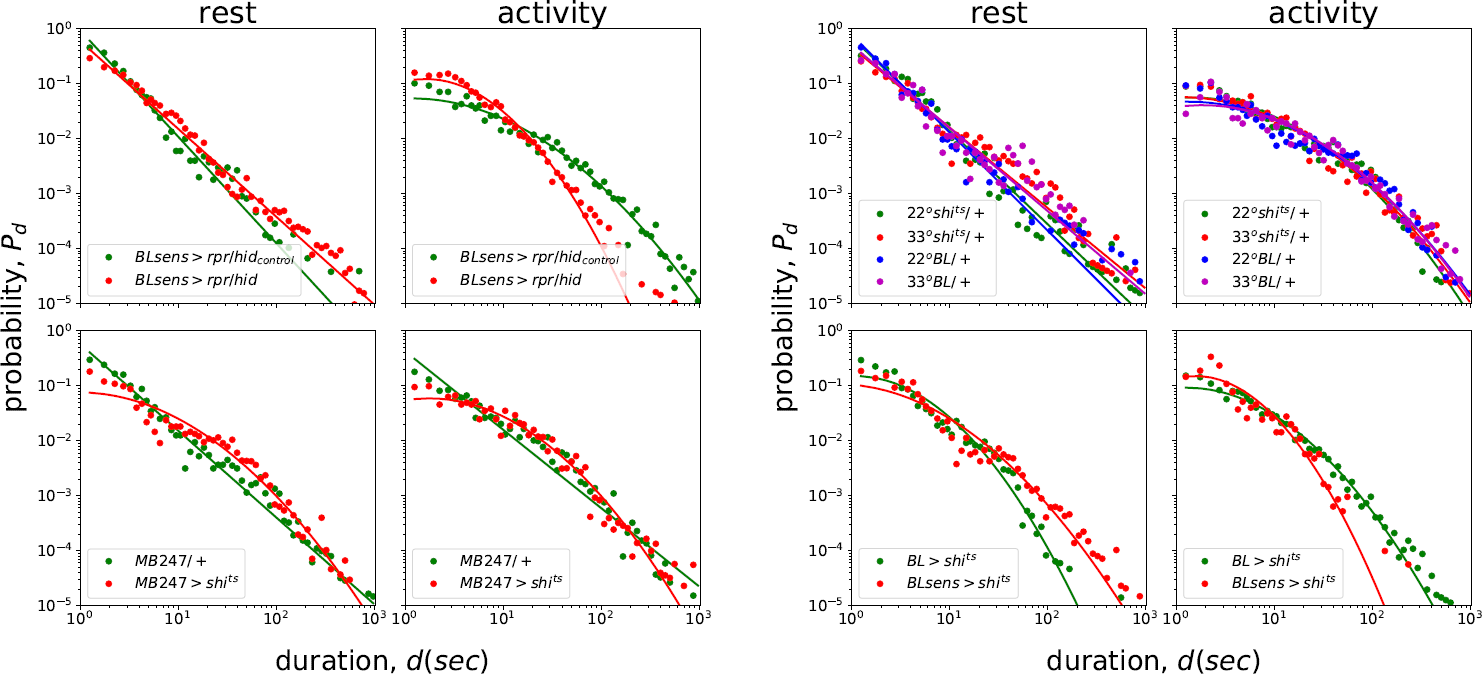
Probability density of rest and activity bout durations for control and activated mutant genotypes. In the first two diagram pairs mutants are plotted against their single respective controls. In the fourth pair the rest two mutants are plotted with their 4 control groups shown in the third diagram pair. Dots describe the probability density over logarithmic bins. Lines indicate the distribution with the lowest Kolmogorov-Smirnov distance *D*_*KS*_ among the best fitting power-law, exponential and log-normal distributions for each group (Tab. 2 - 3).

Interestingly, genetic intervention can have a large effect on the activity ratio. When inactivating sensory neurons and to a lesser extend the mushroom body the activity ratio is decreased (cf. BLsens > *rpr/hid*, MB247 > *shi*^*ts*^ and Blsens > *shi*^*ts*^ in Tab. 1). Inspection of the empirical duration distribution of rest bouts in Fig. 4 (first and third columns) shows that while the power-law fit is superior for all control groups, the log-normal fit approximates best the respective mutant distribution in 3 out of 4 cases (cf. MB247 > *shi*^*ts*^, BL > *shi*^*ts*^ and BLsens > *shi*^*ts*^ in Tab. 2. This might hint impairment of the power-law generating processes due to neural dysfunction. In the fourth case of BLsens > *rpr/hid* the power-law is preserved but shifted to higher values. Regarding activity, the empirical distributions indicate that overall the activity epochs are severely shortened in time for both the BLsens > *rpr/hid* and the BLsens > *shi*^*ts*^ mutants in comparison to the respective control groups (second and fourth columns) hinting early termination of activity bouts by the intermittency mechanism

## Discussion

As most neuroscientific research focuses either on static network connectivity or on neural activation/inhibition - behavior correlations, an integrative account of how temporal behavioral statistical patterns arise from unperturbed neural dynamics is still lacking. In this context, we hope to contribute to scientific discovery in a dual way. Firstly by extending existing mechanistic hypothesis for larva intermittent behavior and secondly by promoting the integration of intermittency in functional models of larval behavior. In what follows we elaborate on these goals and finally describe certain limitations of our study.

### Self-limiting inhibitory waves might underlie intermittent crawling and its modulation

The neural mechanisms underlying intermittency in larva behavior remain partly unknown. Displacement runs are intrinsically discretized, comprised of repetitive, stereotypical peristaltic strides. These stem from segmental central pattern generator circuits (CPG) located in the ventral nerve chord, involving both excitatory and inhibitory premotor neurons and oscillating independently of sensory feedback (7). A ‘visceral pistoning’ mechanism involving head and tail-segment synchronous contraction underlies stride initiation (8). Speed is mainly controlled via stride frequency (8).Crawling is intermittently stopped during both stimulus-free exploratory behavior and chemotaxis, giving rise to non-stereotypical stationary bouts during which reorientation might occur. During the former they are intrinsically generated without need for sensory feedback or brain input (3), while during the latter an olfactory-driven sensorimotor pathway facilitates cessation of runs when navigating down-gradient. Specifically, inhibition of a posterior-segment premotor network by a sub-esophageal zone descending neuron deterministically terminates runs allowing easier reorientation (9).

It is reasonable to assume that this intermittent crawling inhibition is underlying both free exploration and chemotaxis, potentially in the form of transient inhibitory bursts. A neural network controlling the CPG through generation of self-limiting inhibitory waves is well suited for such a role. In the simplest case, during stimulus-free exploration, the durations of the generated inhibitory waves should follow a power-law distribution, behaviorally observed as rest bouts. In contrast, non-power-law distributed quiescent periods of the network would disinhibit locomotion allowing the CPG to generate repetitive peristaltic strides resulting in behaviorally observed runs.

The model we presented (cf. 3.1) alternates autonomously between avalanches of power-law distributed durations and quiescence intervals of exponentially distributed durations without need for external input. Therefore it can serve as a theoretical basis for the development of both generative models that reproduce the intermittent behavior of individual larvae and of the above mechanistic hypothesis for the initiation and cessation of peristaltic locomotion in the larva through disinhibition and inhibition of the crawling CPG respectively. To uncover the underlying neural mechanism and confirm/reject our hypothesis, inhibitory input to the crawling CPG should be sought, measured neurophysiologically and correlated to behaviorally observed stride and stride-free bouts during stimulus-free exploration.

Intermittent behavior in the *Drosophila* adult is subject to two modes of modulation, neither of which affects the distribution of the activity bouts. Firstly, high ambient temperature and daylight raise the activity ratio over long timescales by raising the number of activity bouts (5). This is achieved by lowering the probability of the extremely long rest bouts, without affecting the power-law exponent of the distribution, which coincides with fewer sleep events (> 5 minutes) observed during the day. This modulation is long-lasting and could result from a different constant tonic activation of the system. Secondly, dopamine neuron activation raises the activity ratio acutely by modulation of the power-law exponent upwards (5) skewing locomotion towards the brownian limit. This modulation could be transient in the context of salient phasic stimulation by the environment.

As mentioned above, during chemotaxis larvae perform more and sharpest reorientations, terminating runs when navigating down-gradient. In case the above hold for the larval nervous system as well, a hypothesis integrating both experimental findings could be that this behavior stems from transient olfactory-driven dopaminergically-modulated inhibition of the crawling CPG. Our conceptual model can be extended to address the above claims by adding tonic and/or phasic input.

### Intermittency can extend functional models of larva locomotion

Traditional random walk models fail to capture the temporal dynamics of animal exploration (1). Even when time is taken into account in terms of movement speed, reorientations are assumed to occur acutely. Integrating intermittency can address this limitation allowing for more accurate functional models of autonomous behaving agents. Such virtual agents can then be used in simulations of behavioral experiments promoting neuroscientifically informed hypothesis that advance over current knowledge and generate predictions that can stimulate further empirical work (10).

It is widely assumed that *Drosophila* larva exploration can be descibed as a random walk of discrete non-overlapping runs and reorientations/head-casts (3) or alternatively that it is generated by the concurrent combined activity of a crawler and a turner module generating repetitive oscillatory forward peristaltic strides and lateral bending motions respectively and possibly involving energy transfer between the two mechanical modes (11–13). Both models can easily be upgraded by adding crawling intermittency which might or might not be independent of the lateral bending mechanism. In the discrete-mode case, intermittency can simply control the duration and transitions between runs and head-casts or introduce a third mode of immobile pauses resulting in a temporally unfolding random walk. In the overlapping-mode case the two modules are complemented by a controlling intermittency module forming an interacting triplet. Depending on the crawler-turner interaction and the effect of intermittency on the turner module, multiple locomotory patterns emerge including straight runs, curved runs, stationary head-casts and immobile pauses. This simple extension would allow temporal fitting of generative models to experimental observations in addition to the primarily pursued spatial-trajectory fitting, facilitating the use of calibrated virtual larvae in simulations of behavioral experiments.

### Limitations

A limitation of our study is that due to the singles-pinepoint tracking, it is impossible to determine whether micro-movements occur during the designated inactivity periods, an issue also unclear for adult fruitflies in (5). It follows that in our analysed dataset, immobile pauses, feeding motions and stationary head casts are indistinguishable. Therefore, what we define as rest bouts should be considered as periods lacking at least peristaltic strides but not any locomotory activity. Our relatively low velocity threshold *V*_*θ*_ = 0.085*mm/sec* though allows strict detection of rest bouts as it is evident from the high activity ratio (higher than 0.7 in most control groups). To tackle this, trackings of higher spatial resolution with more spinepoints tracked per larva are needed, despite the computational challenge of the essentially long recording duration.

Also, our results show that an exponential distribution of activity bouts (5) as reported for the adult fruitfly might not be the case for the larva, as we detected log-normal long-tails in all cases. Still, the exponential-power-law duality in our model illustrates switching between independent and coupled modes of neural activity. Substituting the exponential regime by other long-tailed distribution such as log-normal might require assuming more complex interactions between the switching regimes and will be pursued in the future so that generative models of the data can be fit.

## ACKNOWLEDGMENTS

This project was supported by the Research Training Group ‘Neural Circuit Analysis’ (DFG-RTG 1960, grant no. 233886668) and the Research Unit ‘Structure, Plasticity and Behavioral Function of the *Drosophila* mushroom body’(DFG-FOR 2705, grant no. 403329959), funded by the German Research Foundation. M.A. was funded by the UPV/EHU post-doctoral training program ESPDOC17/17 and H2020 Marie Skłodowska-Curie grant 892715, and supported in part from the Basque Government (IT1228-19). We also thank Dr. Jimena Berni for providing an updated version of the larva-tracking dataset, on our request.

The authors do not declare any conflicts of interest.

